# Thiol and Disulfide Containing Vancomycin Derivatives Against Bacterial Resistance and Biofilm Formation

**DOI:** 10.1101/2021.08.23.457343

**Authors:** Inga S. Shchelik, Karl Gademann

## Abstract

Antibiotic-resistant and biofilm-associated infections constitute a rapidly growing issue. Use of the last resort antibiotic vancomycin is under threat, due to the increasing appearance of vancomycin resistant bacteria as well as the formation of biofilms. Herein, we report a series of novel vancomycin derivatives carrying thiol- and disulfide-containing moieties. The new compounds exhibited enhanced antibacterial activity against a broad range of bacterial strains, including vancomycin resistant microbes and Gram-negative bacteria. Moreover, all obtained derivatives demonstrated improved antibiofilm formation activity against VanB resistant *Enterococcus* compared to vancomycin. This work established a promising strategy for combating drug-resistant bacterial infections or disrupting biofilm formation and advances the knowledge on structural optimization of antibiotics with sulfur-containing modifications.

---

Growing multidrug bacterial resistance has become a life-threatening problem.^1^ A fraction of antibiotic-resistant bacteria dubbed “the ESKAPE pathogens” including *Enterococcus faecium* and *Staphylococcus aureus* are listed as high priority targets for new treatments.^2,3^ Several glycopeptide antibiotics have contributed to their sustained effectiveness in the clinic including “last resort” antibiotic vancomycin broadly used against methicillin-resistant *S. aureus* (MRSA). ^4–7^ However, after over 60-years of clinical use, vancomycin-resistant *S. aureus* (VRSA) and vancomycin-resistant *Enterococci* (VRE) have brought new challenges to antibacterial treatment.^8–10^ Consequently, there is a pressing need for renewed antibiotic discovery which overcomes the forces of evolution and selection responsible for bacterial resistance. By avoiding the molecular basis of the prevalent mechanism of resistance and providing additional mechanisms of action, drug potency against the development of resistance can be improved. Vancomycin binds to the C-terminus D-alanyl-D-alanine moiety of bacterial cell wall precursors and inhibits cell wall biosynthesis.^11,12^ The mechanism of resistance to vancomycin involves the alteration of the peptidoglycan synthesis pathway leading to the incorporation of either D-alanyl-D-lactate (VanA, VanB VRE)^13,14^ or D-alanyl-D-serine (VanC VRE)^15^ sequences outside of the cytoplasm. These variations result in the loss of affinity between vancomycin and the peptide and in decreased activity of antibiotic to the resistant strains. Various modification strategies have already been applied to increase vancomycin activity, such as ligand-binding enhancement,^16–18^ binding to the pyrophosphate group of lipid II peptidoglycan,^19,20^ or attachment of bacterial membrane disrupting fragments.^18,21–25^ These approaches involve increasing of the residence time of antibiotics at the bacterial cell therefore improving their performance. In this respect, disulfide units might be considered as an alternative method for retaining the antibiotic on the surface of bacteria or facilitating its uptake. A number of antibiotics contain disulfide units, which have been postulated to be involved in structural rigidification (e.g., in peptide antibiotics)^26–30^ or part of the pharmacophore,^31–42^ show a broad range in activity against both Gram-positive and Gram-negative strains as well as biofilm formation. However, utilizing thiol-mediated uptake in bacteria was rarely investigated, and the results obtained have been mixed so far.^43^

Additionally, to growing resistance, about 80% of all chronic infections are associated with an increased survive ability of a causative pathogen in a formed biofilm.^44–46^ Among them, Gram-positive *cocci* are well established bacteria related to biofilm infections. *Staphylococci* are the leaders of prosthetic related infections^47,48^ followed by *Enterococci*.^49^ Treatment of these infections is not always successful due to bacterial tolerance to conventional antimicrobial agents. The combined effects of the emergence of antibiotic-resistant strains and their ability to form biofilms represent a serious threat nowadays. Therefore, the development of drugs capable of combating both resistant strains and biofilm formation is particularly attractive.

In this work, we investigated the influence of sulfur-modified vancomycin derivatives on antibacterial activity against a broad range of bacterial strains., thereby rationally probing different substitution effects on the sulfur unit. Firstly, a series of vancomycin derivatives were synthesized by C-terminal modification of the glycopeptide with corresponding amines in the presence of PyBOP and HOBt as coupling reagents (Scheme 1). Lipoic acid derivatives (**1a-c**) have been chosen along the line of successful eukaryotic, thiol-mediated uptake systems,^50^ which have earlier also been evaluated with mixed success against bacteria.^43^ Endolipidic disulfides (**2a-c**) were selected to increase lipophilicity, which should also positively influence residence time and potentially involve other mechanisms of actions, such as disruption of cell-wall integrity and inhibition of transglycosylases.^51^ Terminal thiols were selected, both with internal disulfides (**3a/3b**) and without (**4a-d**), which would be most accessible to any bacterial counterpart. Finally, disulfides with polar end groups such as ammonium salts (**5a-c**), pyridine (6) and amine (**7**) were chosen to increase interaction with negatively charged compounds on the bacterial cell wall and improve bacteria membrane permeability.

**Scheme 1.**
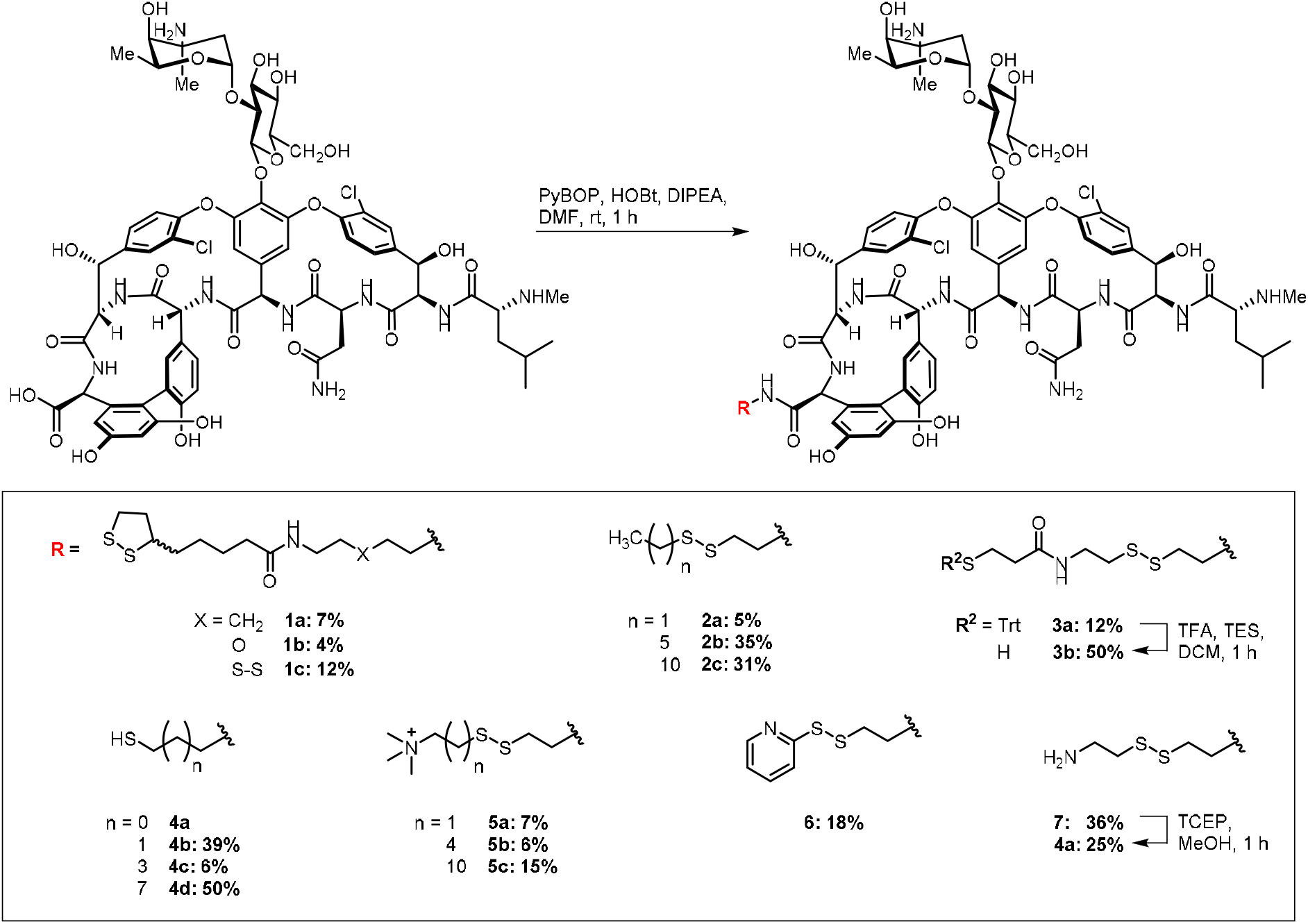
Functionalization of vancomycin at carboxylic position with sulfur-containing linkers.

The antibacterial activities of vancomycin thiol- and disulfide-containing derivatives against a selection of pathogenic clinically isolated Gram-positive strains including resistant MRSA, VISA, and VRE were determined by standard broth microdilution susceptibility tests.^52,53^ The data are summarized in Table 1 (more details in Table S2 in the Supporting Information). The results reveal that the presence of the lipophilic chains is important for antibacterial activity against vancomycin intermediate, vancomycin resistant strains and *S. pneumonia*. The C6 and C11 carbon chains connected to the disulfide derivatives **2b-2c** displayed 16 to 128-fold enhanced activity for VISA, VanA, and VanB strains compared to vancomycin. The derivatives with trimethyl ammonium moiety **5a-5c** were especially active against VISA and *B. subtilis* (Table S2 in the Supporting Information) and demonstrated 8-fold improvement against VRE compared to vancomycin. Moreover, C2-C5 carbon chain derivatives **5a-5b** also showed high increase in activity for the *C. difficile* strain. The enhanced VRE antimicrobial activities of the vancomycin derivatives possessing lipophilic chains (**2b**-**2c**), or quaternary ammonium group (**5c**) might be explained by feasible contribution of the additional mechanism of action associated with disrupting bacterial cell membrane integrity. A similar effect has been reported earlier for the C-terminal modifications of vancomycin with aliphatic chains and trimethylammonium salt.^23,24,54^ The lipophilic C5 chain derivative with a free thiol group **4c** was the most effective antibacterial agent against vancomycin resistant strains and also demonstrated over 16-fold improvement in activity for *S. pneumonia* and *C. difficile*. One reason for these observations might be possible compound dimerization during the *in vitro* experiments. A few examples of disulfide-containing dimers were earlier investigated by Nicolaou et al.^55^ and Sundram et al.^56^ demonstrating an improved activity against resistant *Enterococci*. The vancomycin derivatives with a lipoic acid (**1a-1c**) substitution did not alter the antibacterial profile of parent vancomycin against sensitive or resistant strains, however 4 to 8-fold increase in activity was observed against VISA, *S. pneumoniae* and *C. difficile* for the compound **1a**. The derivatives containing both disulfide bond and terminal thiol group (**3a**) revealed good potential against vancomycin intermediate and resistant strains with 8 to 64-fold enhancement in activity compared to vancomycin. Interestingly, the Trt protected analog (**3b**) of the compound **3a** did not show the loss in activity compared to its deprotected version and furthermore demonstrated the slight improvement (4-fold) against certain *C. difficile* strains and *B. subtilis* (Table S2 in the Supporting Information).

**Table 1.** Minimum Inhibitory Concentrations (MIC) of vancomycin derivatives against selected strains.

|  | MIC (μg/mL) |  |  |  |  |  |  |  |  |  |
| --- | --- | --- | --- | --- | --- | --- | --- | --- | --- | --- |
|  | Sensitive | MRSA | VISA | hVISA | VanA | VanB | VanC |  |  |  |
|  | *S. aureus*<sup>a</sup> | *S. aureus*<sup>b</sup> | *S. aureus*<sup>c</sup> | *S. aureus*<sup>d</sup> | *E. faecium*<sup>e</sup> | *E. faecalis*<sup>f</sup> | *E. gallinarum*<sup>g</sup> | *S. pneumoniae*<sup>h</sup> | *C. difficile*<sup>i</sup> | *M. catarrhalis*<sup>j</sup> |
| **VAN** | 1-2 | 1-2 | 8 | 16-32 | >128 | 16 | 8-16 | 0.12-0.5 | 0.12-0.5 | >32 |
| **1a** | 2 | 2 | 1 | 32 | >32 | 4 | 8 | 0.06 | 0.06 | 4 |
| **1b** | 4 | 2 | n.d. | n.d. | n.d. | 4 | 8 | n.d. | n.d. | n.d. |
| **1c** | 2 | 2 | 2 | 2 | >32 | 4 | 8 | 0.25 | 0.5 | 8 |
| **2a** | 1 | 1 | n.d. | n.d. | n.d. | 4-8 | 8 | n.d. | n.d. | n.d. |
| **2b** | 0.12-0.5 | 0.12-0.25 | 0.5 | 0.5 | 16 | 0.5 | 4-8 | 0.06 | 0.5 | 2 |
| **2c** | 1 | 1-2 | 0.5 | 1 | 2 | 2 | 4 | 0.25 | 0.12 | 0.5 |
| **3a** | 8 | 4 | n.d. | n.d. | n.d. | 2 | 1-2 | n.d. | n.d. | n.d. |
| **3b** | 0.5 | 0.5-1 | 1 | 1 | 8 | 2-4 | 4 | 0.25 | 1 | 2 |
| **4a** | 4 | 4 | n.d. | n.d. | n.d. | 2-4 | 2 | n.d. | n.d. | n.d. |
| **4b** | 4 | 4 | 8 | >32 | 1 | 4 | 1-2 | ≤0.03 | 0.12 | 16 |
| **4c** | 8 | 8 | 2 | >32 | 1 | 1-2 | 0.5-1 | ≤0.03 | 0.06 | 4 |
| **4d** | 4 | 4 | 2 | 2 | 2 | 8 | 8 | 0.5 | 4 | 8 |
| **5a** | 1 | 0.5 | 0.5 | >32 | 32 | 16 | 4 | 0.25 | ≤0.015 | 2 |
| **5b** | 1 | 0.5-1 | 1 | >32 | 16 | 8-16 | 4 | 0.06 | ≤0.015 | 2 |
| **5c** | 1 | 0.5 | 0.25 | 8 | 16 | 2 | 2 | 0.12 | 1 | 1 |
| **6** | 4 | 2-4 | 4 | 4 | 8 | 16 | 8-16 | 0.5 | 0.5 | >32 |
| **7** | 1-2 | 1-2 | 1 | 1 | 8 | 8 | 8 | ≤0.03 | 0.5 | >32 |

VAN: vancomycin, MRSA: methicillin resistant; VISA: vancomycin-intermediate; hVISA: heterogenous vancomycin-intermediate; VanA, VanB, VanC: vancomycin resistant. <sup>a</sup>ATCC 25923, <sup>b</sup>ATCC 43300, <sup>c</sup>ATCC NRS1, <sup>d</sup>ATCC NRS24, <sup>e</sup>MMX0485, <sup>f</sup>ATCC 51299, <sup>g</sup>ATCC 700221, <sup>h</sup>ATCC 29212, <sup>i</sup>ATCC 43255 (rt 087), <sup>j</sup>ATCC 8193. n.d.: not determined. Color coding is relative to VAN activity for each strain separately.

**Table 3.**
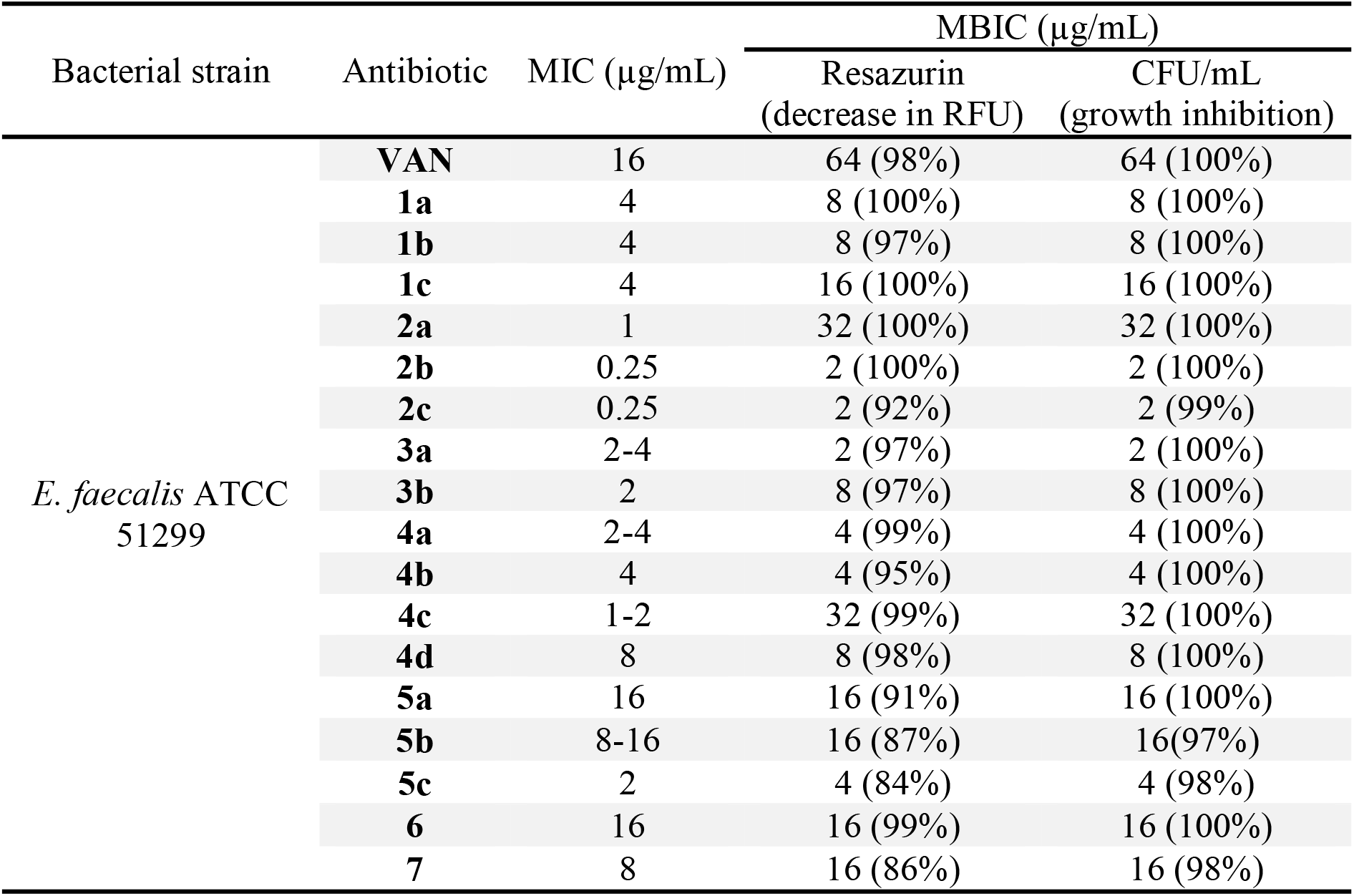
Minimal biofilm inhibitory concentration (MBIC) determined by resazurin and cell counting methods

The lipo-ammonium complex^57^ or sulfonium modification^24^ in vancomycin derivatives were reported to have antibacterial activity against Gram-negative strains. Therefore, we sought to investigate whether the introduction of sulfur-containing linkers could achieve a similar effect. The derivatives were tested against several Gram-negative bacteria (Table S2 in the Supporting Information). Unfortunately, the modifications made did not have any effect on antibacterial activity against *E. coli, A. baumannii* or *P. aeruginosa*. However, we observed high activity (MIC = 1-2 µg/mL) against *M. catarrhalis* strains for the compounds possessing lipophilic chains (**2b-2c**) or Trt group (**3b**) as well as derivatives with quaternary ammonium salt (**5a-5c**) compared to vancomycin, which is resistant to these strains (Table 1). Collectively, these data provide clear and strong evidence that the presence of sulfur-containing linkers enhance the antibacterial profile of vancomycin improving the activity against a broader spectrum of bacterial strains.

Before investigating the inhibitory activity of our compounds on biofilm formation, the optimization of conditions for biofilm growth and the assessment of metabolic activity were firstly carried out. The biofilm-forming activity of six Gram-positive clinically isolated bacterial strains was investigated in six different media: MHB, TS, TSG, TS2G, BHI, and BHIG, after 24 h incubation for *Staphylococci* and *Bacillus*, and 48 h for *Enterococci*.^58^ The assessment of biofilm production carried out by crystal violet (CV) staining and analysis. From the obtained results, the biomass as measured by the absorbance of CV at 570 nm revealed low values for *Enterococci* (lower 0.3) in all tested media (Figure S1 in the Supporting Information). Exceptionally, *E. faecalis* strain had the higher biofilm mass formation, displaying absorbance values ranging from 1.5 to 4.9 in BHI medium supplemented with 1% glucose. *B. subtilis* showed relatively high absorption values in TSG broth, however the biofilm was not uniformly formed therefore was not used for the further experiments. *Staphylococcus* spp. exhibited the highest biofilm-forming profile in BHI broth with values over 3 and over 2 for vancomycin-sensitive and MRSA strains, respectively. Moreover, it was observed that the supplementation of BHI medium with 1% glucose improved the biofilm formation for *Staphylococcus* by 20%. The results of biofilm assessment for the best performers are summarized in Figure 1.

**Figure. 1.**
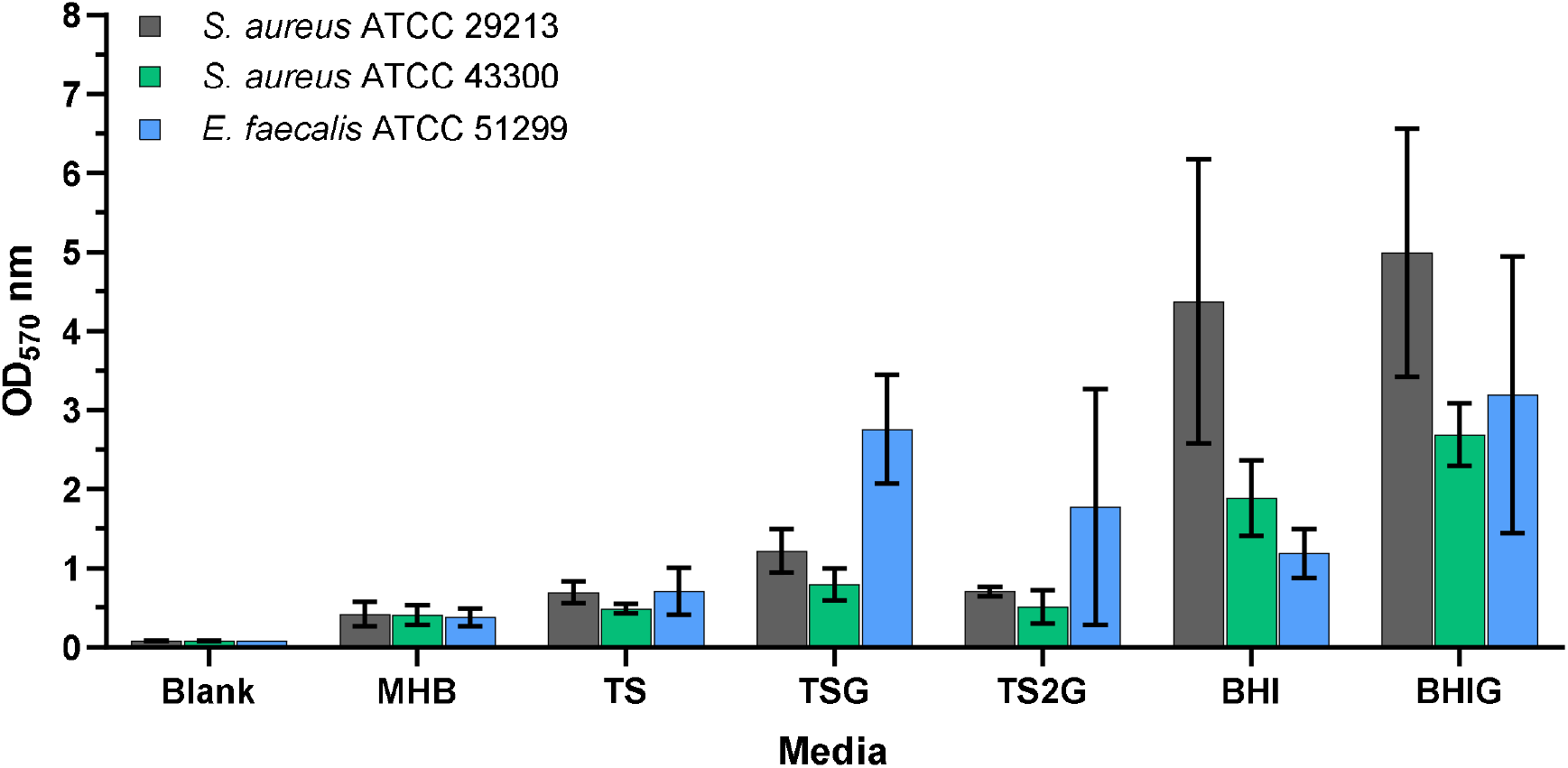
Assessment of biofilm biomass by crystal violet staining of three bacterial strains: *S. aureus* ATCC 29213, *S. aureus* ATCC 43300 (MRSA), *E. faecalis* ATCC 51299. Strains were incubated in six different media. Error bars represent standard deviation of two independent experiments in triplicate.

Composition of the medium is likely the most important factor influencing the ability of bacteria to produce biofilm under *in vitro* conditions. Earlier studies have shown that liquid media (TS or BHI) supplemented with glucose (0.25-4%) initiate greater biofilm formation for staphylococcal and enterococcal species.^58–62^ In our findings, the BHI media supplemented with 1% glucose had the best performance for the *Staphylococci* and *E. faecalis*. Moreover, Mohamed et al. reported that independent of the media, *E. faecalis* produces biofilm stronger than *E. faecium*, which is in agreement with our results.^49^ These results demonstrate the important role of media in biofilm production amongst Gram-positive strains.

Next, we moved to the investigation of metabolic activity of formed biofilms. The previous studies have shown the comparison characteristics between several assays evaluating viable cells and concluded that resazurin is a good alternative to other assessments of metabolic activity of Gram-positive strains.^63^ Therefore, for the assessment of metabolic activity of biofilm cells, the resazurin viability assay was chosen for the Gram-positive strains. Biofilms from the selected strains were formed over 24 h for *S. aureus* and 48 h for *E. faecalis* before the resazurin assay. The formed biofilms were treated with resazurin solutions at concentrations of 2, 4, and 8 µg/mL, respectively, and the plates were incubated at 37 °C. The relative fluorescence units (RFU) were taken every 2 min for 2 h. For all tested strains, the RFU values were increasing during 80 min of measure and then remain constant till the maximum incubation time (Figure S2 in the Supporting Information). The selection of optimal assay conditions was based on calculated statistical quality parameters (Table S5 in the Supporting Info). The conditions chosen for the resazurin assay mostly yielded an excellent Z’ value (≥ 0.5). According to the obtained results the conditions for the resazurin assay selected for further experiments were set as follows: 8 µg/mL with incubation for 60 min for *S. aureus* ATCC 29213, and 4 µg/mL with incubation for 40 min for *S. aureus* ATCC 43300 and *E. faecalis* ATCC 51299.

Using the previously optimized assay, novel synthesized vancomycin derivatives were tested for their ability to repress the growth and biofilm formation by three bacterial strains. The biofilms were grown in optimized earlier conditions in the presence of antibiotics at different concentrations. After incubation, MBIC was determined by resazurin assay with the chosen conditions. Colony forming units (CFU) were also defined in order to corroborate the results obtained with the resazurin-based assay.

The vancomycin derivatives presented MBIC values similar to their MIC data when tested against *Staphylococcus* strains. No significant improvement was observed after the vancomycin modifications compared to vancomycin itself (Table S6 in the Supporting Information). However, all new synthesized compounds completely suppressed biofilm formation of vancomycin-resistant *E. faecalis* at concentrations lower than vancomycin itself (Table 2). In particular, vancomycin derivatives with C6 and C11 lipophilic chains (**2b** and **2c**) as well as containing both a disulfide bond and free thiol group (**3a**) displayed a 32-fold increase in MBIC values compared to vancomycin. Moreover, 16-fold enhancement in inhibition activity was observed for the compounds with shorter aliphatic chains and free thiol group at the end (**4a, 4b**) or compound with longer chain but containing trimethyl ammonium salt (**5c**). Furthermore, when compared to the CFU results, the obtained data revealed the quality and accurateness of these anti-biofilm assays for MBIC determination.

In earlier work by the group of Matile, it was demonstrated that the introduction of cyclic disulfides or selenides into vancomycin structure leads to either no improvement in activity or its slight loss when tested against Gram-positive *B. subtilis*.^43^ Moreover, the Gram-negative strain did not show any sensitivity to the antibiotic after its modification. However, in our work, we showed significant increase of the antibiotic activity against several sensitive and resistant strains including Gram-negative *M. catarrhalis* after the incorporation of either cyclic disulfides or other sulfur-containing linkers into vancomycin structure. The obtained results revealed the improved delivery of the modified antibiotic to several Gram-positive and Gram-negative bacteria, which prompts the further need to study the effect of thiol groups on antibiotics activity or potential thiol-mediated uptake. The high improvement in activity for the vancomycin derivatives with quaternary ammonium salts bearing a terminal tetradecyl substituents was demonstrated by the Boger group when tested against VanA resistant *Enteroccocus*.^23^ Our study revealed that the introduction of a disulfide bond instead of ammonium salt with a long aliphatic chain not only has a similar increase in activity against VanA resistant strains but also appeared to be active against Gram-negative bacteria. Moreover, the combination of both disulfide bond and ammonium salt moieties for the vancomycin modification added activity against *C. difficile*. The nature of high activity and possible dimerization for vancomycin derivatives with free thiol groups should be further investigated. Nevertheless, the introduction of free thiols and disulfide moieties are expected to potentially improve the ADME profile of new antibiotics as a result of metabolization into more hydrophilic compounds for better excretion as was shown in the studies of Mu and colleagues.^64^ Moreover, the modification of the C-terminal position on vancomycin giving the prospect for the further modification of glucosamine part by the introduction of a CBP-group in order to further increase the activity of the antibiotic against vancomycin-resistant strains.^23,65^

In conclusion, we present the potential to improve the antibacterial and antibiofilm formation activity of vancomycin by its structure optimization, with sulfur-containing linkers. This strategy paves the way to new semisynthetic vancomycin analogs for the treatment of vancomycin-resistant bacterial infections and Gram-negative bacteria as well as inhibition of biofilm formation for VanB resistant enterococcus. Overall, this work provides prospective options for further investigation of disulfide and thiol - containing compounds previously underestimated in medicinal chemistry.

## ASSOCIATED CONTENT

The Supporting Information is available free of charge on the ACS Publications website. Supplementary Figures S1-S2, Tables S2, S5, S6, experimental procedures and methods, characterization data, ^1^H and ^13^C NMR spectra, supplementary MIC tables.

## AUTHOR INFORMATION

### Authors

Inga S. Shchelik − Department of Chemistry, University of Zurich, 8057 Zu□rich, Switzerland; orcid.org/0000-0002-0469-683X

### Author Contributions

I. S. S. and K. G. designed the study, I. S. S. performed all experiments, I. S. S. and K. G. discussed the data, I. S. S. and K. G. wrote the manuscript.

## ACKNOWLEDGMENT

The authors acknowledge the Swiss National Science Foundation (SNSF, Grant No. 182043), and a Bundesstipendium for financial support. The authors acknowledge the NMR and Mass spectrometry facilities, the Center for Microscopy and Image Analysis (ZMB) of the University of Zurich for training and maintenance of the instruments.

## ABBREVIATIONS

MIC: minimum inhibitory concentration
MBIC: minimum biofilm inhibitory concentration
CFU: colony forming unit
RFU: relative fluorescent unit

## TOC Graphic

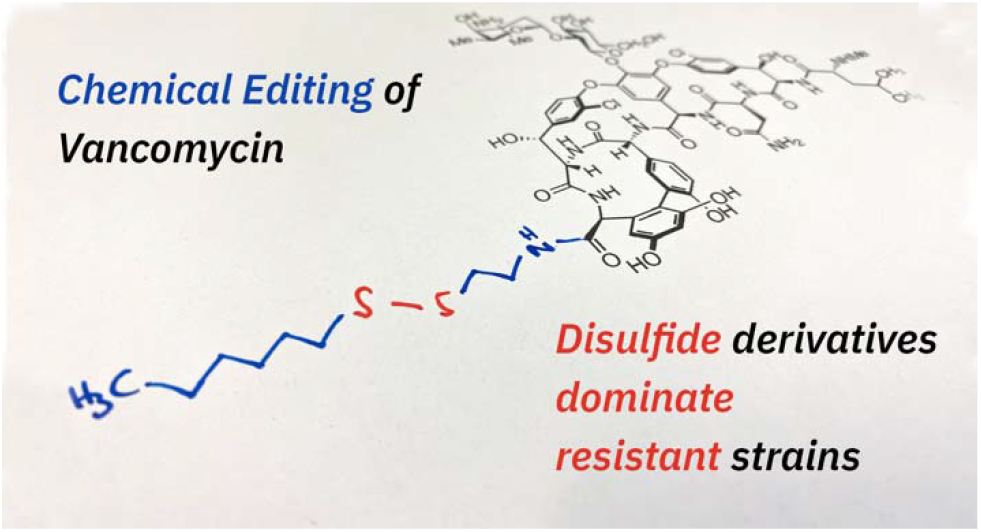

## REFERENCES

(1) Biggest Threats and Data | Antibiotic/Antimicrobial Resistance | CDC https://www.cdc.gov/drugresistance/biggest-threats.html (accessed 2021-08-23).

(2) Willyard, C. The Drug-Resistant Bacteria That Pose the Greatest Health Threats. Nature, 2017, 543, 15.

(3) Pendleton, J. N.,, Gorman, S. P.,, Gilmore, B. F. Clinical Relevance of the ESKAPE Pathogens. Expert Rev. Anti-Infect. Ther. 2013, 3, 297–308.

(4) Kahne, D.; Leimkuhler, C.; Lu, W.; Walsh, C. Glycopeptide and Lipoglycopeptide Antibiotics. Chem. Rev. 2005, 2, 425–448.

(5) Blaskovich, M. A. T.; Hansford, K. A.; Butler, M. S.; Jia, Z.; Mark, A. E.; Cooper, M. A. Developments in Glycopeptide Antibiotics. ACS Inf. Dis. 2018, 4, 715–735.

(6) McCormick, M. H.; Stark, W. M.; Pittenger, G. E.; Pittenger, R. C.; McGuire, J. M. Vancomycin, a New Antibiotic. I: Chemical and Biologic Properties. Antibiot. Annu. 1955, 1, 606–611.

(7) Nicolaou, K. C.; Boddy, C. N. C.; Bräse, S.; Winssinger, N. Chemistry, Biology, and Medicine of the Glycopeptide Antibiotics. Angew. Chem. Int. Ed. 1999, 38, 2096–2152.

(8) Walsh, C. Vancomycin Resistance: Decoding the Molecular Logic. Science 1993, 261, 5119.

(9) Mcguinness, W. A.; Malachowa, N.; Deleo, F. R. Vancomycin Resistance in Staphylococcus aureus. Yale J. Biol. Med. 2017, 90, 269–281.

(10) Miller, W. R.; Munita, J. M.; Arias, C. A. Mechanisms of Antibiotic Resistance in Enterococci. Expert Rev. Anti-infect. Ther. 2014, 12, 1221–1236.

(11) Barna, J. C. J.; Williams, D. H. The Structure and Mode of Action of Glycopeptide Antibiotics of the Vancomycin Group. Ann. Rev. Microbiol. 1984, 38, 339–357.

(12) Perkins, H. R. Vancomycin and Related Antibiotics. Pharmacol. Ther. 1982, 16, 181–197.

(13) Handwerger, S., Pucci M. J.; Volk, K. J.; Liu, J.; Lee, A. S. The Cytoplasmic Peptidoglycan Precursor of Vancomycin Resistant Enterococcus faecalis Terminates in Lactate. J. Bacteriol. 1992, 174, 5982–5984.

(14) Evers, S.; Courvalin, P. Regulation of VanB-Type Vancomycin Resistance Gene Expression by the VanS_B_-VanR_B_ Two-Component Regulatory System in Enterococcus faecalis V583. J. Bacteriol. 1996, 178, 1302–1309.

(15) S. Dutka-Malen, C. Molinas, M. Arthur, P. Courvalin, Sequence of the vanC gene of Enterococcus gallinarum BM4174 encoding a D-alanine:D-alanine ligase-related protein necessary for vancomycin resistance. Gene 1992, 112, 53–58.

(16) Yarlagadda, V.; Konai, M. M.; Manjunath, G. B.; Ghosh, C.; Haldar, J. Tackling Vancomycin-Resistant Bacteria with “Lipophilic-Vancomycin-Carbohydrate Conjugates.” J. Antibiot. 2015, 68, 302–312.

(17) Guan, D.; Chen, F.; Xiong, L.; Tang, F.; Faridoon Qiu, Y.; Zhang, N.; Gong, L.; Li, J.; Lan, L.; Huang, W. Extra Sugar on Vancomycin: New Analogues for Combating Multidrug-Resistant Staphylococcus Aureus and Vancomycin-Resistant Enterococci. J. Med. Chem. 2018, 61, 286–304.

(18) Yarlagadda, V.; Samaddar, S.; Paramanandham, K.; Shome, B. R.; Haldar, J. Membrane Disruption and Enhanced Inhibition of Cell-Wall Biosynthesis: A Synergistic Approach to Tackle Vancomycin-Resistant Bacteria. Angew. Chem. Int. Ed. 2015, 54 (46), 13644– 13649.

(19) Yarlagadda, V.; Sarkar, P.; Samaddar, S.; Haldar, J. A Vancomycin Derivative with a Pyrophosphate-Binding Group: A Strategy to Combat Vancomycin-Resistant Bacteria. Angew. Chem. Int. Ed. 2016, 55, 7836–7840.

(20) Guan, D.; Chen, F.; Faridoon Liu, J.; Li, J.; Lan, L.; Huang, W. Design and Synthesis of Pyrophosphate-Targeting Vancomycin Derivatives for Combating Vancomycin-Resistant Enterococci. ChemMedChem 2018, 13, 1644–1657.

(21) Yarlagadda, V.; Akkapeddi, P.; Manjunath, G. B.; Haldar, J.Membrane Active Vancomycin Analogues: A Strategy to Combat Bacterial Resistance. J. Med. Chem. 2014, 57, 4558–4568.

(22) Blaskovich, M. A. T.; Hansford, K. A.; Gong, Y.; Butler, M. S.; Muldoon, C.; Huang, J. X.; Ramu, S.; Silva, A. B.; Cheng, M.; Kavanagh, A. M.; Ziora, Z.; Premraj, R.; Lindahl, F.; Bradford, T. A.; Lee, J. C.; Karoli, T.; Pelingon, R.; Edwards, D. J.; Amado, M.; Elliott, A. G.; Phetsang, W.; Daud, N. H.; Deecke, J. E.; Sidjabat, H. E.; Ramaologa, S.; Zuegg, J.; Betley, J. R.; Beevers, A. P. G.; Smith, R. A. G.; Roberts, J. A.; Paterson, D. L.; Cooper, M. A. Protein-Inspired Antibiotics Active against Vancomycin-and Daptomycin-Resistant Bacteria. Nat. Comm. 2018, 9, 1–17.

(23) Okano, A.; Isley, N. A.; Boger, D. L. Peripheral Modifications of [Ψ[CH_2_NH]Tpg4]Vancomycin with Added Synergistic Mechanisms of Action Provide Durable and Potent Antibiotics. Proc. Natl. Acad. Sci. U. S. A. 2017, 114, E5052–E5061.

(24) Guan, D.; Chen, F.; Qiu, Y.; Jiang, B.; Gong, L.; Lan, L.; Huang, W. Sulfonium, an Underestimated Moiety for Structural Modification, Alters the Antibacterial Profile of Vancomycin Against Multidrug□Resistant Bacteria. Angew. Chem. Int. Ed. 2019, 58, 6678–6682.

(25) Yarlagadda, V.; Sarkar, P.; Manjunath, G. B.; Haldar, J. Lipophilic Vancomycin Aglycon Dimer with High Activity against Vancomycin-Resistant Bacteria. Bioorg. Med. Chem. Lett. 2015, 25, 5477–5480.

(26) Vetterli, S. U.; Zerbe, K.; Müller, M.; Urfer, M.; Mondal, M.; Wang, S.-Y.; Moehle, K.; Zerbe, O.; Vitale, A.; Pessi, G.; Eberl, L.; Wollscheid, B.; Robinson, J. A. Thanatin Targets the Intermembrane Protein Complex Required for Lipopolysaccharide Transport in Escherichia coli. Sci. Adv. 2018, 4, eaau2634.

(27) Essig, A.; Hofmann, D.; Münch, D.; Gayathri, S.; Künzler, M.; Kallio, P. T.; Sahl, H. G.; Wider, G.; Schneider, T. Aebi, M. Copsin, a Novel Peptide-Based Fungal Antibiotic Interfering with the Peptidoglycan Synthesis. J. Biol. Chem. 2014, 289, 34953–34964.

(28) Bédard, F.; Hammami, R.; Zirah, S.; Rebuffat, S.; Fliss, I.; Biron, E. Synthesis, Antimicrobial Activity and Conformational Analysis of the Class IIa Bacteriocin Pediocin PA-1 and Analogs Thereof. Sci. Rep. 2018, 8, 9029.

(29) Duprez, W.; Premkumar, L.; Halili, M. A.; Lindahl, F.; Reid, R. C.; Fairlie, D. P.; Martin, J. L. Peptide Inhibitors of the Escherichia Coli DsbA Oxidative Machinery Essential for Bacterial Virulence. J. Med. Chem. 2015, 58, 577–587.

(30) Rabanal, F.; Grau-Campistany, A.; Vila-Farrés, X.; Gonzalez-Linares, J.; Borràs, M.; Vila, J.; Manresa, A.; Cajal, Y. A Bioinspired Peptide Scaffold with High Antibiotic Activity and Low in Vivo Toxicity. Sci. Rep. 2015, 5, 10558.

(31) Cavallito, C. J.; Bailey, J. H. Allicin, the Antibacterial Principle of Allium Sativum. I. Isolation, Physical Properties and Antibacterial Action. J. Am. Chem. Soc. 1944, 66, 1950–1951.

(32) Tsao, S. -M.; Yin, M. -C. In Vitro Activity of Garlic Oil and Four Diallyl Sulphides against Antibiotic-Resistant Pseudomonas Aeruginosa and Klebsiella Pneumoniae. J. Antimicrob. Chem. 2001, 47, 665–670.

(33) Jakobsen, T. H.; van Gennip, M.; Phipps, R. K.; Shanmugham, M. S.; Christensen, L. D.; Alhede, M.; Skindersoe, M. E.; Rasmussen, T. B.; Friedrich, K.; Uthe, F.; Jensen, P. Ø.; Moser, C.; Nielsen, K. F.; Eberl, L.; Larsen, T. O.; Tanner, D.; Høiby, N.; Bjarnsholt, T.; Givskov, M. Ajoene, a Sulfur-Rich Molecule from Garlic, Inhibits Genes Controlled by Quorum Sensing. Antimicrob. Agents Chemother. 2012, 56, 2314–2325.

(34) Kim, D.; Sun Lee, I.; Hyung Jung, J.; Yang, S.-I. Psammaplin A, a Natural Bromotyrosine Derivative from a Sponge, Possesses the Antibacterial Activity against Methicillin-Resistant Staphylococcus aureus and the DNA Gyrase-Inhibitory Activity. 1999, 22, 25–29.

(35) Nicolaou, K. C.; Hughes, R.; Pfefferkorn, J. A.; Barluenga, S. Optimization and Mechanistic Studies of Psammaplin A Type Antibacterial Agents Active against Methicillin-Resistant Staphylococcus aureus (MRSA). Chem. Eur. J. 2001, 7, 4296–4310.

(36) Brinkhoff, T.; Bach, G.; Heidorn, T.; Liang, L.; Schlingloff, A.; Simon, M. Antibiotic Production by a Roseobacter Clade-Affiliated Species from the German Wadden Sea and Its Antagonistic Effects on Indigenous Isolates. Appl. Environ. Microbiol. 2004, 70, 2560– 2565.

(37) Beyersmann, P. G.; Tomasch, J.; Son, K.; Stocker, R.; Göker, M.; Wagner-Döbler, I.; Simon, M.; Brinkhoff, T. Dual Function of Tropodithietic Acid as Antibiotic and Signaling Molecule in Global Gene Regulation of the Probiotic Bacterium Phaeobacter Inhibens. Sci. Rep. 2017, 7, 730.

(38) Wright W.G.; Watson K.C. Antibiotic activity of Gerrardine and Cassipourine, alkaloids from plants of the family Rhizophoraceae. S. Afr. Med. J. 1967, 94–96.

(39) Chan, A. N.; Shiver, A. L.; Wever, W. J.; Razvi, S. Z. A.; Traxler, M. F.; Li, B. Role for Dithiolopyrrolones in Disrupting Bacterial Metal Homeostasis. Proc. Natl. Acad. Sci. 2017, 114, 2717–2722.

(40) Sheppard, J. G.; Frazier, K. R.; Saralkar, P.; Hossain, M. F.; Geldenhuys, W. J.; Long, T. E. Disulfiram-Based Disulfides as Narrow-Spectrum Antibacterial Agents. Bioorg. Med. Chem. Lett. 2018, 28, 1298–1302.

(41) Danquah, C. A.; Kakagianni, E.; Khondkar, P.; Maitra, A.; Rahman, M.; Evangelopoulos, D.; McHugh, T. D.; Stapleton, P.; Malkinson, J.; Bhakta, S.; Gibbons, S. Analogues of Disulfides from Allium Stipitatum Demonstrate Potent Anti-Tubercular Activities through Drug Efflux Pump and Biofilm Inhibition. Sci. Rep. 2018, 8, 1150.

(42) Sakai, K.; Iwatsuki, M.; Iizuka, M.; Asami, Y.; Nonaka, K.; Masuma, R.; Takizawa, M.; Nakashima, T.; Tokiwa, T.; Shiomi, K.; Ōmura, S. Aldsulfin, a Novel Unusual Anti-Mannheimiosis Epithiodiketopiperazine Antibiotic Produced by Lasiodiplodia Pseudotheobromae FKI-4499. J. Antibiot. 2021, 74, 363–369.

(43) Laurent, Q.; Berthet, M.; Cheng, Y.; Sakai, N.; Barluenga, S.; Winssinger, N.; Matile, S. Probing for Thiol-Mediated Uptake into Bacteria. ChemBioChem 2020, 21, 69–73.

(44) del Pozo, J. L. R. P. The Challenge of Treating Biofilm-Associated Bacterial Infections. Clin. Pharmacol. Ther. 2007, 82, 204–209.

(45) Lynch, A. S.; Robertson, G. T. Bacterial and Fungal Biofilm Infections. Annu. Rev. Med. 2008, 59, 415–428.

(46) Jamal, M.; Ahmad, W.; Andleeb, S.; Jalil, F.; Imran, M.; Nawaz, M. A.; Hussain, T.; Ali, M.; Rafiq, M.; Kamil, M. A. Bacterial Biofilm and Associated Infections. J. Chin. Med. Assoc. 2018, 81, 7–11.

(47) Chu, V. H.; Crosslin, D. R.; Friedman, J. Y.; Reed, S. D.; Cabell, C. H.; Griffiths, R. I.; Masselink, L. E.; Kaye, K. S.; Corey, G. R.; Reller, L. B.; Stryjewski, M. E.; Schulman, K. A.; Fowler, V. G. Staphylococcus aureus Bacteremia in Patients with Prosthetic Devices: Costs and Outcomes. Am. J. Med. 2005, 118, 1416.e19-1416.e24.

(48) Costerton, J. W.; Stewart, P. S.; Greenberg, E. P. Bacterial Biofilms: A Common Cause of Persistent Infections. Science, 1999, 284, 1318–1322.

(49) Mohamed, J. A.; Huang, D. B. Biofilm Formation by Enterococci. J. Med. Microbiol. 2007, 56, 1581–1588.

(50) Laurent, Q.; Martinent, R.; Lim, B.; Pham, A.-T.; Kato, T.; López-Andarias, J.; Sakai, N.; Matile, S. Thiol-Mediated Uptake. JACS Au, 2021, 1, 710–728.

(51) Ge, M.; Chen, Z.; Russell, H.; Onishi Kohler, J.; Silver, L. L.; Kerns, R.; Fukuzawa, S.; Thompson, C.; Kahne, D. Vancomycin Derivatives That Inhibit Peptidoglycan Biosynthesis Without Binding D-Ala-D-Ala. Science 1999, 284, 507–511.

(52) Abmm, D.; Tamma, D.; Kirn, J.; Cullen, S. K. Clinical and Laboratory Standards Institute (CLSI). Performance Standards for Antimicrobial Susceptibility Testing. 30th Ed. CLSI supplement M100 2020, Wayne, PA.

(53) EUCAST Reading Guide for Broth Microdilution; 2021.

(54) Wu, Z.-C.; Boger, D. L. Exploration of the Site-Specific Nature and Generalizability of a Trimethylammonium Salt Modification on Vancomycin: A-Ring Derivatives. Tetrahedron 2019, 75, 3160–3165.

(55) Nicolaou, K. C.; Hughes, R.; Cho, S. Y.; Winssinger, N.; Labischinski, H.; Endermann, R. Synthesis and Biological Evaluation of Vancomycin Dimers with Potent Activity against Vancomycin-Resistant Bacteria: Target-Accelerated Combinatorial Synthesis. Chem. Eur. J. 2001, 7 (17), 3824–3843.

(56) Burgess, K.; Zhu, J. Novel Vancomycin Dimers with Activity against Vancomycin-Resistant Enterococci. Chemtracts 1997, 10 (11), 818–821.

(57) Yarlagadda, V.; Manjunath, G. B.; Sarkar, P.; Akkapeddi, P.; Paramanandham, K.; Shome, B. R.; Ravikumar, R.; Haldar, J. Glycopeptide Antibiotic To Overcome the Intrinsic Resistance of Gram-Negative Bacteria. ACS Infectious Diseases 2016, 2 (2). https://doi.org/10.1021/acsinfecdis.5b00114.

(58) Cruz, C. D.; Shah, S.; Tammela, P. Defining Conditions for Biofilm Inhibition and Eradication Assays for Gram-Positive Clinical Reference Strains. BMC Microbiol. 2018, 18, 173.

(59) Knobloch, J. K. M.; Horstkotte, M. A.; Rohde, H.; Mack, D. Evaluation of Different Detection Methods of Biofilm Formation in Staphylococcus aureus. Med. Microbiol. Immunol. 2002, 191, 101–106.

(60) Stepanović, S.; Vuković, D.; Hola, V.; di Bonaventura, G.; Djukić, S.; Ćirković, I.; Ruzicka, F. Quantification of Biofilm in Microtiter Plates: Overview of Testing Conditions and Practical Recommendations for Assessment of Biofilm Production by Staphylococci. APMIS 2007, 115, 891–899.

(61) Pillai, S. K.; Sakoulas, G.; Eliopoulos, G. M.; Moellering, R. C.; Murray, B. E.; Inouye, R. T. Effects of Glucose on Fsr-Mediated Biofilm Formation in Enterococcus Faecalis. J. Inf. Dis. 2004, 190, 967–970.

(62) Ran, S. J.; Jiang, W.; Zhu, C. L.; Liang, J. P. Exploration of the Mechanisms of Biofilm Formation by Enterococcus Faecalis in Glucose Starvation Environments. Aust. Dent. J. 2015, 60, 143–153.

(63) Peeters, E.; Nelis, H. J.; Coenye, T. Comparison of Multiple Methods for Quantification of Microbial Biofilms Grown in Microtiter Plates. J. Microbiol. Methods 2008, 72, 157–165.

(64) Mu, Y.; Nodwell, M.; Pace, J. L.; Shaw, J.-P.; Judice, J. K. Vancomycin Disulfide Derivatives as Antibacterial Agents. Bioorg. Med. Chem. Lett. 2004, 14, 735–738.

(65) Wu, Z.-C.; Isley, N. A.; Boger, D. L. N-Terminus Alkylation of Vancomycin: Ligand Binding Affinity, Antimicrobial Activity, and Site-Specific Nature of Quaternary Trimethylammonium Salt Modification. ACS Inf. Dis. 2018, 4, 1468–1474.

